# Does Adaptive Protein Evolution Proceed by Large or Small Steps at the Amino Acid Level?

**DOI:** 10.1101/379073

**Authors:** Juraj Bergman, Adam Eyre-Walker

## Abstract

A longstanding question in evolutionary biology is the relative contribution of large and small effect mutations to the adaptive process. We have investigated this question in proteins by estimating the rate of adaptive evolution between all pairs of amino acids separated by one mutational step using a McDonald-Kreitman type approach and genome-wide data from several *Drosophila* species. We find that the rate of adaptive evolution is higher amongst amino acids that are more similar. This is partly due to the fact that the proportion of mutations that are adaptive is higher amongst more similar amino acids. We also find that the rate of neutral evolution between amino acids is higher amongst similar amino acids. Overall our results suggest that both the adaptive and non-adaptive evolution of proteins is dominated by substitutions between amino acids that are more similar.

## Introduction

Whether evolution proceeds by large or small steps is an old evolutionary problem that dates back, in its most extreme form, to the debate between saltationists and gradualists at the turn of the 20th century. It is a problem that is far from resolved despite extensive theoretical and experimental work (Barrett and Schluter, 2008, Bell, 2009, Rockman, 2012).

There are in fact three related questions relating to the contribution of large and small mutations to the adaptive process: what is the distribution of effect sizes amongst new mutations, what is the distribution of those that spread to fixation, and is the process of adaptation largely a consequence of large or small mutations. An analogy might help to illustrate the difference between the last two questions. Let us imagine that a builder is constructing a wall. The supply of bricks may be dominated by either large or small bricks and depending on her preferences for bricks three different walls may be built; one in which most of the bricks are small and the wall is largely constructed of small bricks, one in which most of the bricks are small but the wall is largely built of large bricks and one in which most of the bricks are large and the wall is largely composed of large bricks.

Fisher (1930) originally suggested, based on his geometric model, that most advantageous mutations would be of small effect. While some experiments have been consistent with this expectation (Bataillon et al., 2011, Kassen and Bataillon, 2006, Sanjuan et al., 2004, Schenk et al., 2012) others have found a relatively uniform (Ferris et al., 2007, MacLean and Buckling, 2009) or normal distribution of effects (McDonald et al., 2011). The difference between these studies seems to be largely a consequence of two factors; a tendency to under-sample mutations with weak effects because they are difficult to detect and how far the population is from the optimum. The further the population is from the optimum the more large-effect mutations are found (MacLean and Buckling, 2009).

The distribution of mutant effects is however not the distribution of mutations fixed during evolution because large effect mutations have a greater chance of spreading to fixation (Kimura 1983). Theoretical work has suggested that the distribution of effects amongst mutations that spread to fixation is likely to be dominated by mutations of small effect if adaptation comes from new mutations, the underlying distribution of mutant effects is of the Gumbel (e.g. a normal distribution) or Weibull (e.g. a distribution with a truncated right tail) type and the fitness optimum moves suddenly (Martin and Lenormand, 2008, Orr, 1998, Orr, 2002) (though see critique by (Kopp and Hermisson, 2009)). However, if the optimum moves slowly, or most adaptation comes from standing genetic variation (Barrett and Schluter, 2008, Pritchard et al., 2010), then substitutions of intermediate effect are expected to dominate the adaptive process (Kopp and Hermisson, 2009, Matuszewski et al., 2014, Matuszewski et al., 2015). The distribution of substitution effects may be dominated by large effect mutations if the underlying distribution of mutant effects is heavy tailed (i.e. in the Frechet domain) (Seetharaman and Jain, 2014).

Experiments that have tracked mutations that either fix or spread to high frequency under positive selection have found that the distribution can be dominated by mutations of small (Imhof and Schlotterer, 2001, Perfeito et al., 2007) or intermediate effect (Barrett et al., 2006, Rokyta et al., 2005, Rozen et al., 2002, Schoustra et al., 2009). This again seems to depend on how far the population is from the optimum. If the population is far from the optimum, as in the experiment of Barrett et al. (Barrett et al., 2006), then the distribution of mutations that rise to appreciable frequency, or are fixed, is dominated by intermediate or large effect mutations, because the distribution of new mutations is dominated by larger effect mutations (see above) and such mutations have a greater chance of spreading through the population. A second factor also comes into play in experiments, which are usually conducted with asexual organisms – clonal interference. If there is clonal interference, then only mutations with intermediate or large effects can spread to high frequency or fixation (see (Perfeito et al., 2007)).

It is unclear however what these experiments tell us about adaptation in the natural world. All experiments assume that adaptation comes from new genetic variation, but this process might be dominated by standing genetic variation (Barrett and Schluter, 2008, Pritchard et al., 2010). Furthermore, clonal interference occurs in many of the experiments and it is not clear how many organisms are sufficiently asexual for this process to play an important role in adaptation. Finally, we have no idea whether evolution is dominated by large jumps in the optimum, as might be caused by the introduction of an antibiotic or a pesticide into the environment, or more gradual changes. The only experiments that would seem to give us information about what happens in the natural world are QTL analyses of the differences between species. These seem to suggest that much adaptation is due the fixation of mutations of large effect (Bell, 2009), but as Rockman (Rockman, 2012) has argued, some caution must be exercised because a single QTL may involve many mutations of smaller effect.

Here we investigate whether adaptive evolution in proteins is dominated by mutations and substitutions between amino acid that are more or less similar to each other in their physicochemical properties. We expect mutations between similar amino acids to be subject to weaker selection than those between different amino acids, a conjecture we test.

There is indeed some evidence for this. Grantham (Grantham, 1974) and Miyata et al. (Miyata et al., 1979) showed many years ago that the rate of amino acid substitution is negatively correlated to the difference in polarity, volume and chemical composition of the amino acids involved (see also (Zhang, 2000)). This could either be due to mutations between more different amino acids being more deleterious or less advantageous. In our analysis, we estimate the rate of adaptive substitution between all pairs of amino acids separated by one mutational step using polymorphism data from *Drosophila melanogaster* polarized using *D. simulans* and *D. yakuba*. We also investigate whether mutations of large or small effect are more common and whether small or large steps contribute most to the increase in fitness.

## Results

To investigate whether adaptive evolution is dominated by large or small steps at the molecular level we estimated the rate of adaptive evolution between all 75 pairs of amino acids that are separated by a single mutational step. We estimated the rates of substitution between *Drosophila melanogaster* and the *D. simulans/D. yakuba* outgroup pair using the method of Schneider et al. (Schneider et al., 2011). This method is a variant of the McDonald-Kreitman (McDonald and Kreitman, 1991) approach in which the rate of adaptive evolution is estimated by comparing the divergence at selected non-synonymous and neutral synonymous sites, to levels of polymorphisms at those same sites. The method estimates the distribution of fitness effects (DFE) of the neutral and deleterious non-synonymous mutations, the proportion of mutations that are advantageous (*λ*_a_) and the strength of selection acting upon them multiplied by the effective population size (*N*_e_*s*_a_), as well as the rate of adaptive evolution relative to the mutation rate (*ω*_a_) (Gossmann et al., 2010). We initially focus our analysis on two properties of amino acids, volume and polarity, since these are two properties that all amino acids share and that have been studied before (Grantham, 1974, Miyata et al., 1979, Zhang, 2000). However, we also consider other measures of physicochemical and evolutionary amino acid dissimilarity. We consider autosomal and X-linked loci separately since mutations on the X are hemizygous in males and there is some evidence that X-linked genes adapt faster (reviewed by Charlesworth et al. (Charlesworth et al., 2018)).

We find that the rate of adaptive evolution relative to the mutation rate, ω_a_, is significantly negatively correlated to both the difference in volume (Δ_vol_) and polarity (Δ_pol_), on both the autosomes and X-chromosome (Table 1; Figure 1A, B for autosomes; Figure S1A, B for X-chromosome) suggesting that the rate of adaptive evolution is higher between amino acids that are more physicochemically similar. The difference in volume and polarity are only weakly correlated (Spearman’s *ρ* = 0.13, p = 0.11) and the two factors are independently correlated to ω_a_ in a multiple regression (p<0.001 for both factors on the autosomes and X).

**Table 1.**
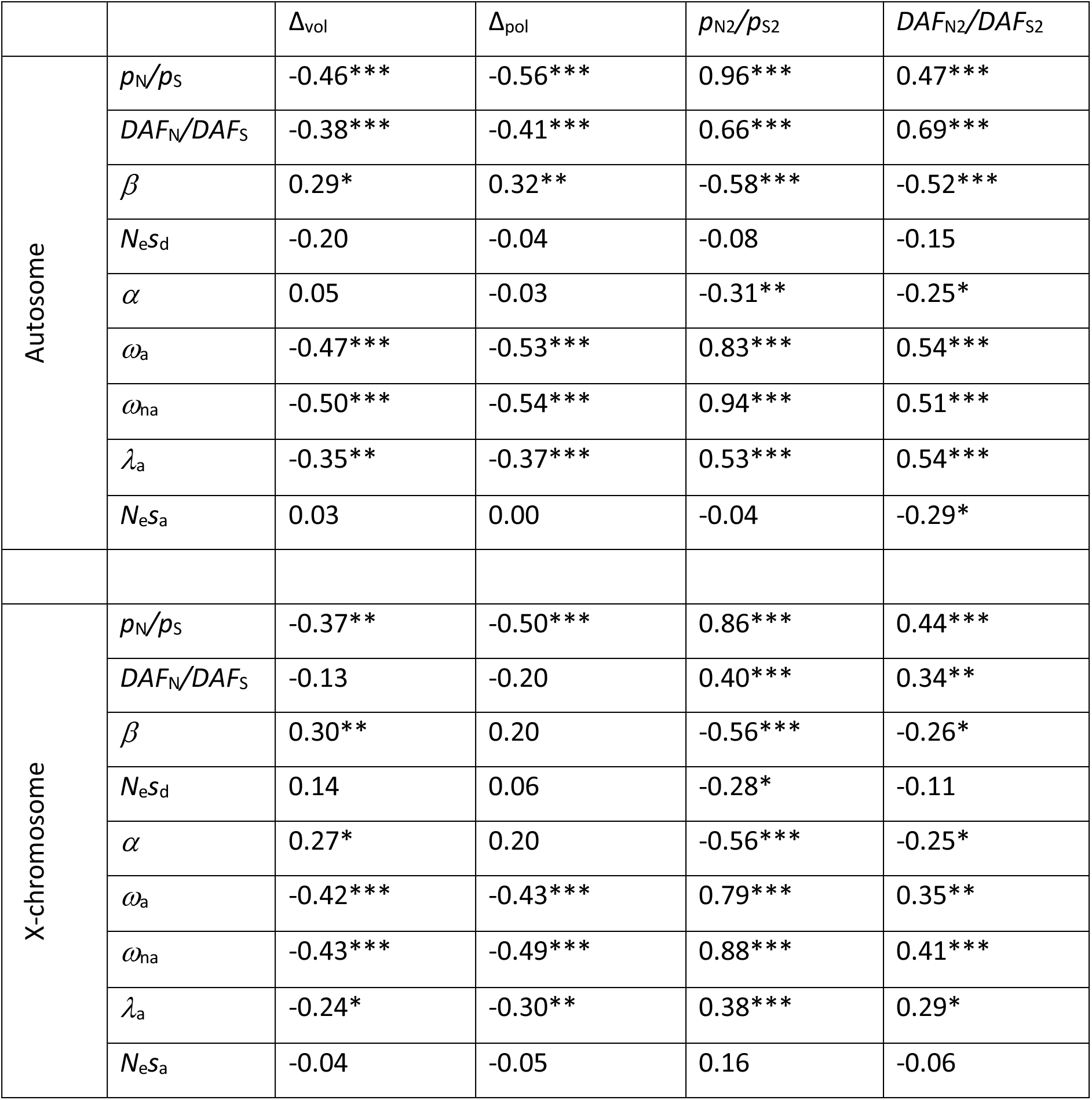
Spearman rank correlation between estimates of rates of adaptive evolution, the DFE, and measures of amino acid dissimilarity. To remove statistical non-independence between *p*_N_/*p*_S_ and *DAF*_N_/*DAF*_S_ and other variables we sampled the SFS to generate two independent SFSs * p < 0.05, ** p < 0.01, *** p < 0.001.

**Figure 1.**
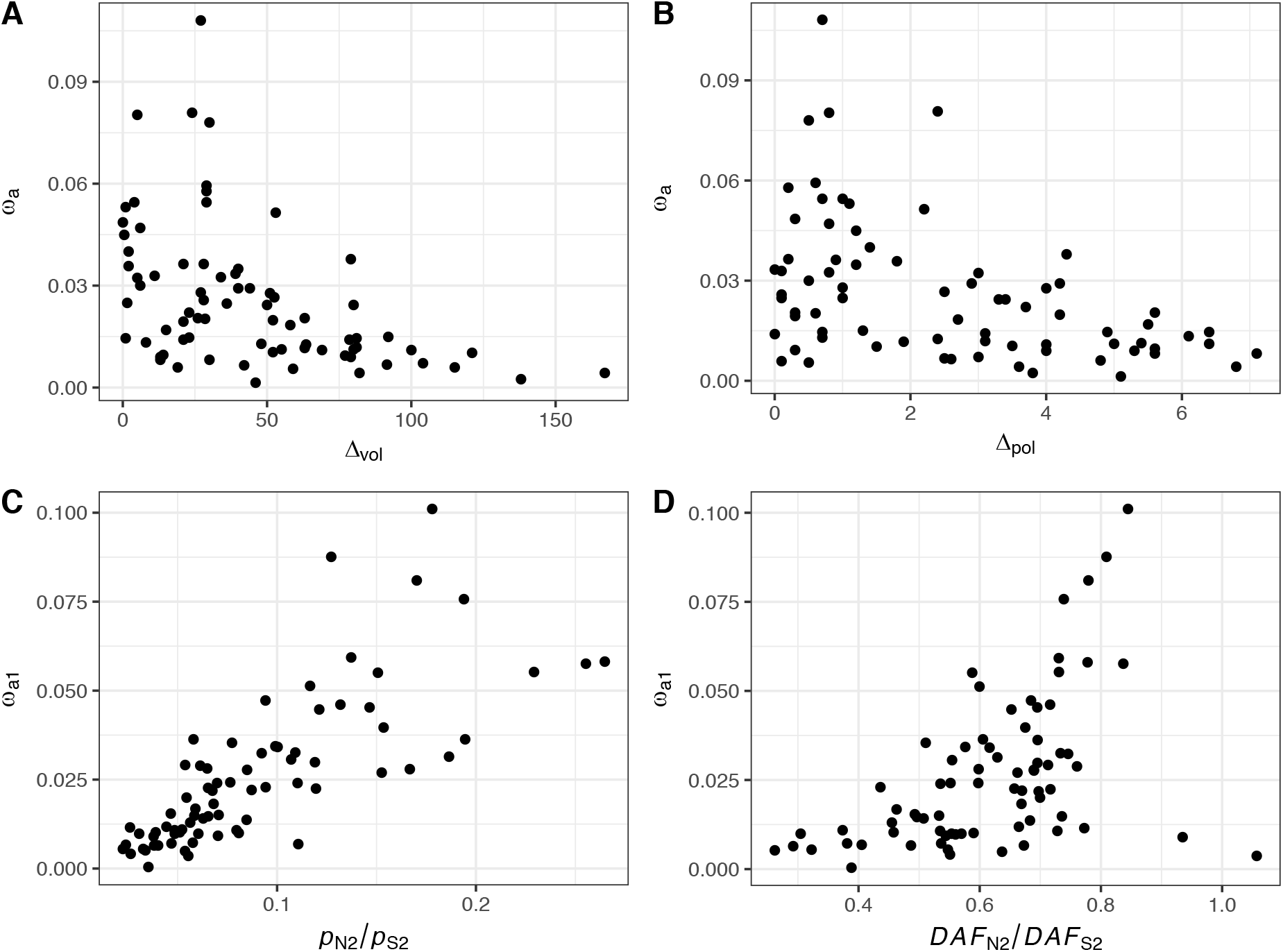
The autosomal rate of adaptive evolution relative to the mutation rate (ω_a_) plotted against the difference in A) volume, B) polarity, C) *p*_N_/*p*_S_ and D) *DAF*_N_/*DAF*_S_.

There are many ways in which to measure the dissimilarity between amino acids, and there are over 500 dissimilarity matrices (Kawashima et al., 2008). We find that *ω*_a_ is negatively correlated to the difference in amino acid properties in ~90% (476/531) of these matrices and significantly so in ~54% (286/531) matrices (Figure 2). *ω*_a_ is positively correlated to the difference in amino acid properties in 55 matrices but none of these correlations are significant.

**Figure 2.**
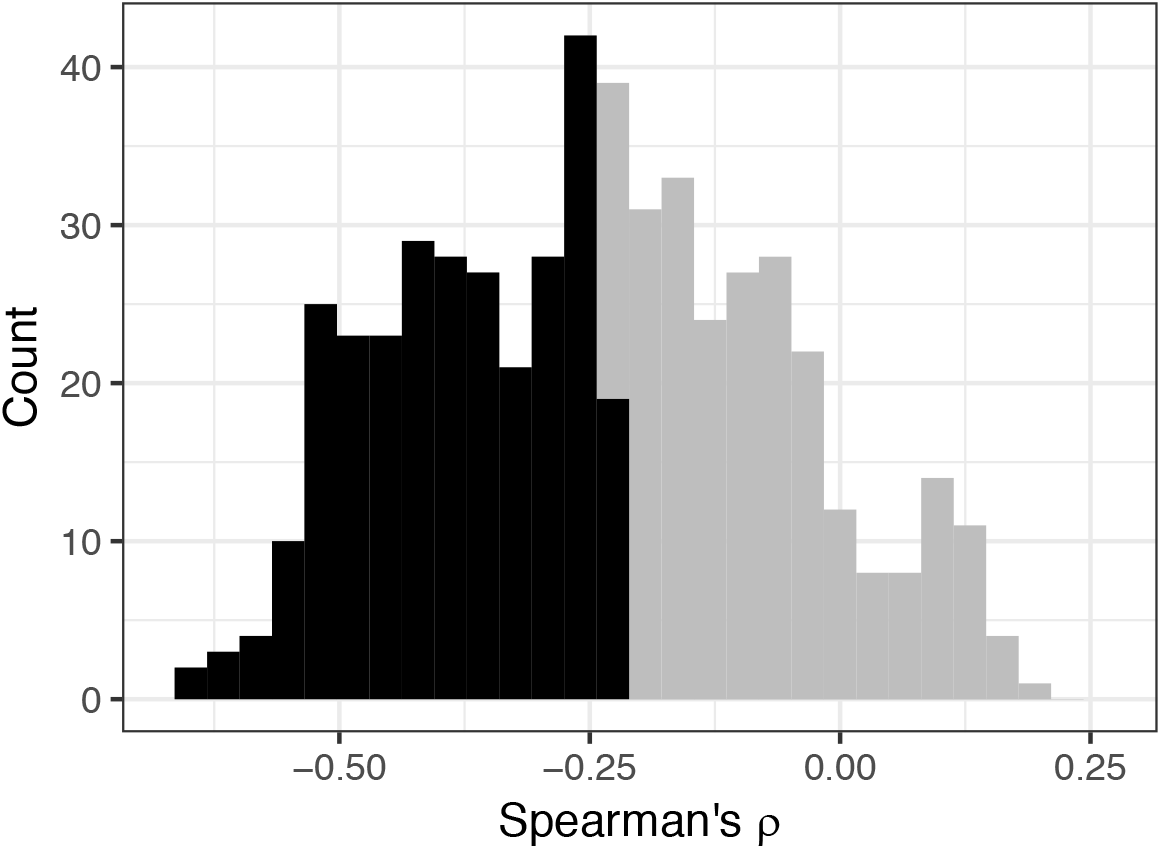
The distribution of Spearman rank correlations between ω_a_ and 531 amino acid dissimilarity matrices. The correlations in the darker shaded area are significant at 5%.

So far we have shown that the rate of adaptive evolution is higher between pairs of amino acids that are more similar in terms of volume and polarity. However, if dissimilar pairs of amino acids tend to be more common or have higher mutation rates, then the overall adaptive evolution might be dominated by substitutions of intermediate or large effect. As a consequence we calculated the total rate of adaptive substitution between each pair of amino acid as Ω_a(ij)_ = ω_a(ij)_ × (*f*_i_ + *f*_j_) × *μ*_ij_ where ω_a(ij)_ is the ω_a_ between a pair of amino acids *i* and *j, f*_i_ is the frequency of amino acid *i* and *μ*_ij_ is the mutation between them; we estimate the mutation rate from synonymous sites (e.g. if the amino acids are separated by a C<>T transition, we estimate the C<>T mutation rate from synonymous sites). If we plot the cumulative number of adaptive amino acids substitutions as a function of the difference in volume and polarity we find the relationship is concave suggesting that small substitutions dominate the adaptive process when we take into account the frequencies and mutation rates of the amino acids (Figure 3A for autosomes; Figure S2A for the X-chromosome).

**Figure 3.**
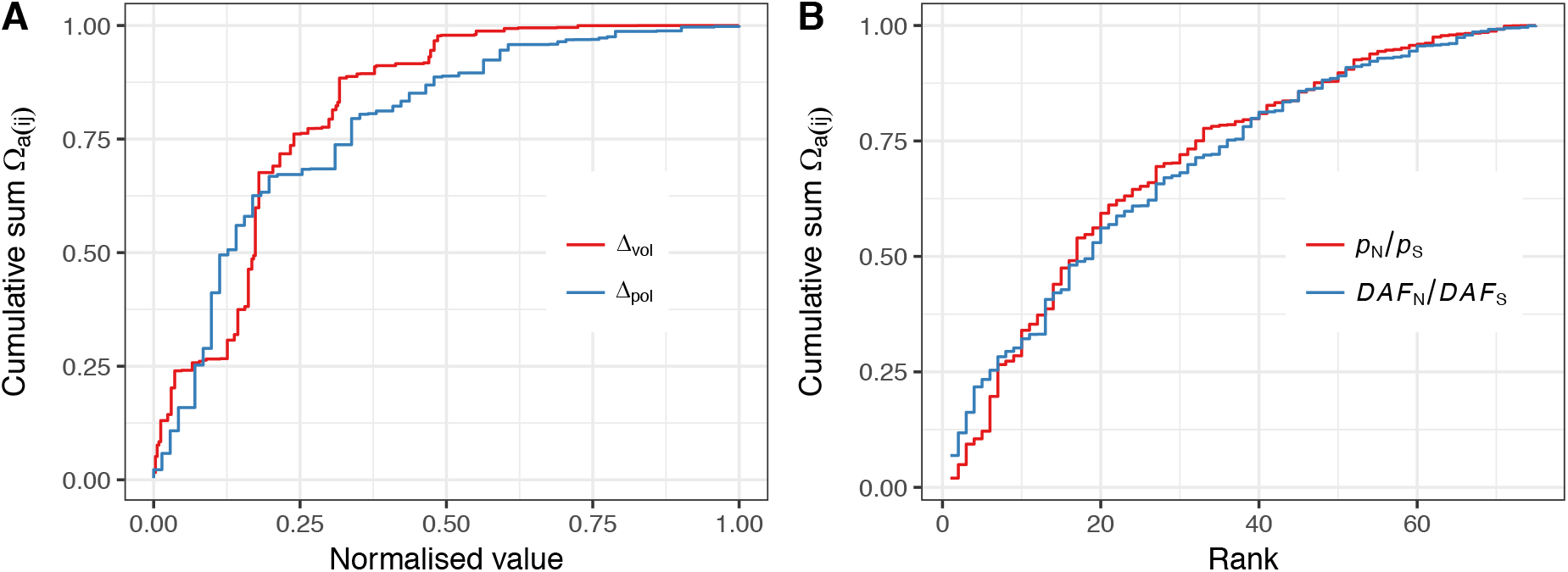
The cumulative number of adaptive substitutions on the autosomes contributed by each pair of amino acids versus A) the normalised difference in volume and polarity, and B) the reverse rank of *p*_N_/*p*_S_ and *DAF*_N_/*DAF*_S_. The normalised difference in volume and polarity was calculated by subtracting the minimum difference, and then dividing by the maximum difference minus the minimum difference.

The fact that more similar amino acids have higher rates of adaptive evolution strongly suggests that the proportion of mutations that are adaptive is also higher amongst more similar amino acids, since more similar amino acids are likely to be subject to weaker selection and hence have lower fixation probabilities. We indeed observe this; *λ*_a_ is significantly negatively correlated to the difference in volume and polarity on both the X and autosomes (Table 1). If we calculate the overall rate of advantageous mutation for each pair of amino acids taking into account the frequency of the amino acids and their mutation rate as Λ_a(ij)_ = *λ*_a(ij)_ × (*f*_i_ + *f*_j_) × *μ*_ij_ and plot the cumulative, we again find that it is concave (Figure S3A, C).

Polarity and volume only explain some of the variance in *ω*_a_ and *λ*_a_, particularly amongst amino acids that are similar in volume or polarity. This is not surprising; volume and polarity are just two measures of amino acid dissimilarity and there are many qualities that are difficult to measure – for example the ability to form disulphide bridges. Alternative measures of amino acid dissimilarity are evolutionary measures such as the ratio of non-synonymous to synonymous polymorphisms (*p*_N_/*p*_S_) and the derived allele frequency of non-synonymous relative to synonymous polymorphisms (*DAF*_N_/*DAF*_S_). Both of these statistics are expected to be higher for amino acids that are more similar because they are expected to decline as the strength of selection against deleterious mutations increases. Consistent with this we find that *p*_N_/*p*_S_ and *DAF*_N_/*DAF*_S_ are negatively correlated to the difference in volume and polarity (Table 1; Figure 4).

**Figure 4.**
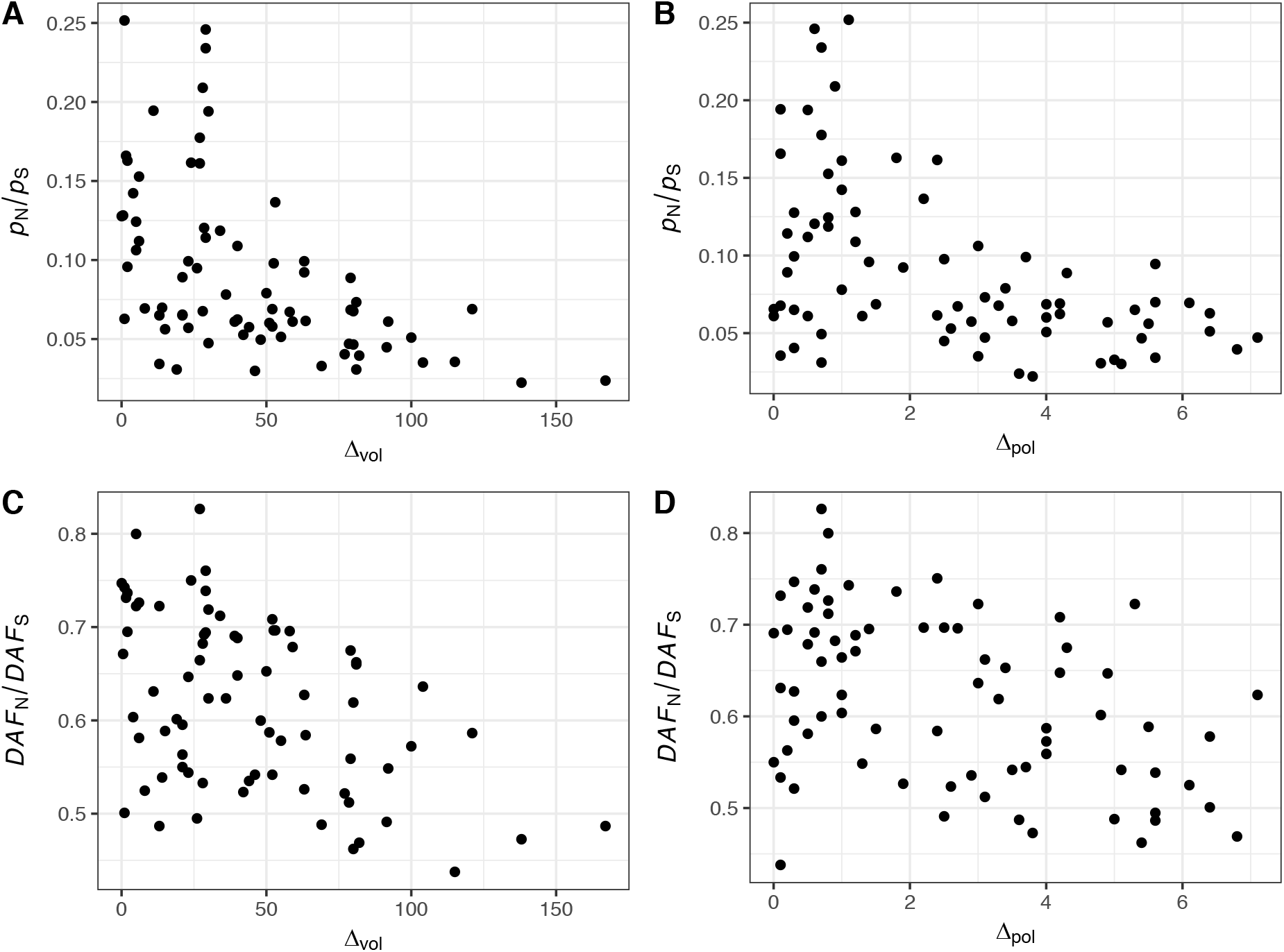
*p*_N_/*p*_S_ and *DAF*_N_/*DAF*_S_ plotted against the difference in volume and polarity for 75 pairs of amino acids for the autosomal data.

Our two evolutionary measures of amino acid dissimilarity, *p*_N_/*p*_S_ and *DAF*_N_/*DAF*_S_ are not statistically independent of our measures of adaptive evolution, since polymorphism data is used to estimate the rate of adaptive evolution; sampling error will therefore tend to induce correlations between ωa, *p*_N_/*p*_S_ and *DAF*_N_/*DAF*_S_. To overcome this, we resampled the SFS using a hypergeometric distribution to generate two SFSs, one of which was used to estimate *p*_N_/*p*_S_ and *DAF*_N_/*DAF*_S_, and the other which was used to estimate the DFE and the rate of adaptive evolution. This procedure removes the non-independence due to sampling error, although we note that *p_N1_*/*p*_S1_ and *p_N2_*/*p*_S2_, are very highly correlated to each other suggesting that there is relatively little sampling error relative to the systematic variance in *p*_N_/*p*_S_ (autosome Spearman’s ρ = 0.96, p<0.001; X-chromosome Spearman’s ρ = 0.86, p<0.001); the correlation between *DAF*_N1_/*DAF*_S1_ and *DAF*_N2_/*DAF*_S2_ is also substantial on the autosomes (autosomes, Spearman’s *ρ* = 0.69, p<0.001; X-chromosome Spearman’s *ρ* = 0.34, p = 0.003). We find that ω_a1_ is significantly positively correlated to *p_N2_*/*p*_S2_ and *DAF*_N2_/*DAF*_S2_ (Table 1, Figure 1C, D). This is consistent with the pattern seen for volume and polarity; amino acids which are more similar in terms of the fitness effects, have high values of *p*_N_/*p*_S_ and *DAF*_N_/*DAF*_S_, and higher rates of adaptive evolution. We also find that the proportion of mutations that are adaptive, *λ*_a1_, is positively correlated to *p_N2_*/*p*_S2_ and *DAF*_N2_/*DAF*_S2_ (Table 1), again consistent with the pattern seen for polarity and volume. If we calculate the overall rates of adaptive substitution, Ω_a(ij)_, and mutation, Λ_a(ij)_ and plot the cumulatives against the ranks of the *p*_N2_/*p*_S2_ and *DAF*_N2_/*DAF*_S2_ values in reverse order, we again observe concave functions (Figure 3B, S2B, S3B, D). Note that we plot the cumulatives against the rank because *p*_N_/*p*_S_ and *DAF*_N_/*DAF*_S_ do not directly relate to any meaningful measure (e.g. they are not simple linear functions of the strength of selection), and we plot them in reverse order because large values correspond to more similar amino acids.

We have shown that both the rate of advantageous mutation and substitution is higher amongst amino acids that are more similar, where we have measured similarity both in terms of physicochemical and evolutionary differences. Finally, we would also like to know whether similar or dissimilar amino acids contribute more overall to adaptation. This question only makes sense phrased in terms of fitness. In principle, we can estimate the contribution of each amino acid pair to the change in fitness by multiplying the rate of adaptive evolution by the mean strength of selection acting on the advantageous substitutions. In principle it is possible to estimate the mean strength of selection from the site frequency spectrum, with or without considering the rate of substitution (Schneider et al., 2011, Tataru et al., 2017). In practice, very large amounts of data are required. We find that our estimate of the strength of selection acting on advantageous mutations is uncorrelated to either the difference in volume, polarity, *p*_N_/*p*_S_ or *DAF*_N_/*DAF*_S_ (Table 1), which suggests that either the strength of selection acting on advantageous mutations is uncorrelated to the similarity of the amino acids, which seems unlikely, or that we cannot estimate the selection strength accurately enough. To assess the sampling error involved in estimating the strength of selection we bootstrapped the data 100 times for the 5 amino acid pairs for which we have the most non-synonymous polymorphisms. Despite having over 1500 non-synonymous polymorphisms in each case we find the confidence intervals span more than one order of magnitude (Figure S4). The reason for this uncertainty is evident upon a visual inspection of the SFSs (Figure S5). Under a model in which non-synonymous mutations are neutral or deleterious the ratio of the non-synonymous and synonymous SFS is expected to be a declining function. However, if there are advantageous mutations the ratio of SFS can be U-shaped and the uptick in the ratio at high allele frequencies contains information about the rate of advantageous mutation and the strength of selection acting upon those mutations (Schneider et al., 2011, Tataru et al., 2017). This signature is subtle and that the ratio of the SFSs is too erratic to infer anything about the strength of selection acting on advantageous mutations (Figure S5).

## Discussion

We have investigated whether the rate of advantageous mutation and substitution depends on the similarity of amino acids. We find that pairs of amino acids that are more similar have higher rates of advantageous mutation and substitution. The adaptive process therefore seems to be dominated by mutations and substitutions of small effect. This is true when we consider the amino acid pairs individually and when we take into account their frequency and mutation rates. However, we have been unable to ascertain whether the overall change in fitness is dominated by small or large mutations. Using the analogy from the introduction, we have established that the supply of bricks is dominated by small bricks and that our builder prefers small bricks for building her wall, so there are more small bricks in the wall. However, we have been unable to establish whether the wall is largely made of small or large bricks.

Our work builds on the work of Grantham (Grantham, 1974) and Miyata et al. (Miyata et al., 1979) who showed, more than 40 years ago, that the rate of evolution is faster between amino acids that are more similar in their physicochemical properties. This might have been because more dissimilar amino acids have lower rates of adaptive evolution, lower rates of neutral evolution or both. We have shown that it is in part due to a lower rate of adaptive evolution (Table 1), but we can also test whether the rate of non-adaptive evolution *ω*_na_ = *d*_N_/*d*_S_ - *ω*_a_ (where *d*_N_ and *d*_S_ are rates of non-synonymous and synonymous divergence, respectively) (Galtier, 2016) is correlated to amino acid dissimilarity. We find that *ω*_na_ is negatively correlated to the difference in volume or polarity, and positively correlated to *p*_N_/*p*_S_ and *DAF*_N_/*DAF*_S_ (Table 1). The fact that both the rate of adaptive and non-adaptive evolution decreases with increasing dissimilarity between amino acids suggests that the proportion of substitutions that are adaptive, *α*, might be relatively constant. We find, however, the proportion of substitutions that are adaptive, *α*, is significantly negatively correlated *p*_N_/*p*_S_ and *DAF*_N_/*DAF*_S_ and significantly positively correlated to the difference in polarity on the X-chromosome (Table 1); i.e. the proportion of substitutions that are adaptive is lower amongst amino acids that are more similar.

Our results may explain the findings of Bazykin and Kondrashov (Bazykin and Kondrashov, 2012) and Campos et al. (Campos et al., 2017). Bazykin and Kondrashov (Bazykin and Kondrashov, 2012) observed that the rate of adaptive amino acid substitution was higher in regions of the gene that were less conserved. Campos et al. (Campos et al., 2017) estimated that the rate of adaptive mutation was lower in more constrained genes, and surprisingly that the strength of selection acting upon those mutations was also weaker. Together these two inferences suggest that constrained genes would also undergo lower rates of adaptive substitution. Hence both analyses mirror at the gene and sub-gene level what we observe at the amino acid level. This begs the question whether genes and parts of genes adapt slowly because of the amino acids they contain, or whether certain amino acids have low rates of adaptive evolution because they tend to be found in genes and parts of genes that have low rates of adaptation. The fact that we observe strong correlations between rates of adaptive evolution at physicochemical properties suggests the former is at least partly true; genes and parts of genes that are constrained undergo low rates of adaptive evolution because they contain amino acids such as glycine which is small, leading to large volume differences, with amino acids that are one mutational step removed from it.

It is striking that much of the variance between amino acids in their rate of adaptive evolution can be explained in terms of *p*_N_/*p*_S_. Given that polymorphism data is expected to be dominated by neutral and slightly deleterious genetic variation, *p*_N_/*p*_S_ is an estimate of the proportion of mutations that are effectively neutral and hence 1-*p*_N_/*p*_S_ a measure of the proportion of mutations that are deleterious. In part the correlation between ω_a_ and *p*_N_/*p*_S_ is not surprising; as amino acids become more different so we expect the proportion of mutations that are effectively neutral to decline, and this is also likely to lead to a reduction in the proportion of mutations that are advantageous, as we have shown (Table 1).

However, we might have also expected advantageous mutations between dissimilar amino acids to be more strongly selected (though see (Campos et al., 2017)). We have been unable to ascertain whether this is the case (*N*_e_*S*_a_ is not significantly correlated to any measure of dissimilarity). However, we can conclude that the strength of selection acting upon advantageous mutations either decreases as amino acid dissimilarity increases or stays constant, neither of which is very likely, or that it increases, but at a low rate, because the rate of adaptive substitution declines as amino acid similarity decreases; i.e. if the strength of selection acting upon advantageous mutations increased rapidly with increasing amino acid dissimilarity then the rate of adaptive evolution would be greater amongst more dissimilar amino acids, even though the proportion of mutations that are adaptive declines as amino acids become more dissimilar.

A potential problem in any analysis that uses the McDonald-Kreitman (MK) approach to estimate the rate of adaptive evolution are differences between the current Ne and the Ne during the divergence phase of evolution, if there is a class of mutations that are slightly deleterious (Eyre-Walker, 2002, McDonald and Kreitman, 1991). If the current Ne, which is relevant to the polymorphism data, is greater than the Ne for the divergence data then MK approaches will tend to overestimate the rate of adaptive evolution; the bias can be such that a signal of adaptive evolution can be detected even when there is no adaptive evolution occurring (Eyre-Walker, 2002, McDonald and Kreitman, 1991). It is not possible for us to rule out this as an explanation for the patterns we observe; the correlation between the rate of adaptive evolution and amino acid similarity might simply be a consequence of increasing population size.

The method that we have used to estimate the rate of adaptive evolution assumes that synonymous mutations are neutral, whereas selection is known to act upon synonymous codon use in some *Drosophila* species (Akashi, 1995, Shields et al., 1988). However, such selection is unlikely to affect our results because the rate of adaptive evolution is estimated using synonymous data that is common to multiple amino acid pairs that are separated by a particular type of mutation (e.g. C<>T). Selection on synonymous codon use could potentially affect the absolute rate of adaptive evolution but it’s not expected to affect the pattern between pairs of amino acids. To investigate further we ran an analysis of covariance regressing *ω*_a_ against the difference in volume and polarity, with mutational type as a fixed effect (in effect fitting a series of parallel planes of *ω*_a_ against the difference in volume and polarity for each mutational type). We find that *ω*_a_ is significantly correlated to the difference in both volume (p<0.001) and polarity (p<0.001). It is also possible that biased gene conversion could affect our results so we repeated the ANCOVA restricting our analysis to GC-conservative mutational types and again find that *ω*_a_ is significantly correlated to the difference in polarity (p<0.001) and volume (p<0.001).

Although we have shown that more similar amino acids undergo higher rates of advantageous mutation and substitution this does not directly address the underlying question of whether adaptive evolution is dominated by small or large effect mutations for two reasons. First, we have only considered amino acid mutations, but much adaptive evolution might proceed through regulatory changes (Andolfatto, 2005, King and Wilson, 1975). Second, underlying each amino acid pair is a distribution of effects; so, although we have shown that the average rate of advantageous mutation and substitution is correlated to measures of amino acid similarity, this does not imply that the underlying distribution, the distribution obtained by combining the distributions from each pair of amino acids, has the same shape. Overall, the adaptive process might be dominated by mutations and substitutions of intermediate effect, but the mean for each of the amino acid distributions is such that they lie to the right of mode of the underlying distribution (Figure S6).

In conclusion, whether evolution proceeds by large or small steps is a long-standing question. We have shown that the adaptation of protein coding sequences is dominated by amino acid mutations that are of small effect.

## Material and methods

### Data and filtering

A population dataset of Zambian *D. melanogaster* sequences was taken from Lack et al. (Lack et al., 2015). In total, the dataset consists of 197 sequences for each autosome and 196 sequences for the X chromosome. Sequences were annotated using the reference genome annotation of *D. melanogaster*} (r5.57 from http://www.flybase.org/) and subsequently masked for all non-coding regions to exclude genomic regions where coding and non-coding sequences overlap. Codon alignments were then extracted using a custom Python script. The alignment between the *D. melanogaster, D. simulans* and *D. yakuba* reference sequences was taken from Hu et al. (Hu et al., 2013). Coding sequences which contained premature stop codons in the *D. melanogaster* reference sequence were excluded from the analysis.

Amino acid polarity scores and volumes were taken from the literature. Additionally, we analyzed other amino acid distance measures using data available in the AAindex1 database (Kawashima et al., 2008). Specifically, for each index in the database, we calculated the physicochemical distance for all amino acid pairs under consideration, as the absolute difference. Indices which contained missing values for any amino acid were excluded from the analysis.

### Parameter inference

We used the method of Schneider et al. (Schneider et al., 2011) to infer the rate of adaptive evolution for all 75 pairs of amino acids separated by a single mutational step. The method requires the unfolded site frequency spectrum (SFS) from a class of sites subject to selection, here non-synonymous sites, and a class of sites in which mutations are neutral, here synonymous sites. Inference of the unfolded site frequency spectrum for each of the site classes was obtained by the method of Keightley et al. (Keightley et al., 2016), using *D. simulans* and *D. yakuba* as outgroups for polarization of *D. melanogaster* sites into ancestral and derived allelic states. Although the dataset contains 197 and 196 lines for the autosomal and X-linked loci, we down-sampled the data to 20 lines. The subsampling step was necessary due to the limited size of the transition matrix used by the program for estimating the demography parameters. Most amino acid pairs are separated by one of the six different mutational types. To estimate the rate of adaptive substitution we compared the SFS for a particular amino acid pair, say proline and threonine, which are separated by a C<>A change with synonymous data from 4-fold degenerate codons separated only by C<>A mutations (SFS_4F(C<>A)_). For amino acids separated by more than one mutational type we calculated a weighted average SFS from the SFSs for the mutational types at 4-fold sites, weighting by the frequency of the respective codons. For example, leucine and valine are separated by C<>G and T<>G. The synonymous SFS used to estimate the rate of adaptive substitution was estimated as SFS_4F(weighted)_ =((*f*_TTA_ + *f*_GTA_ + *f*_TTG_ + *f*_GTG_) × SFS_4F(T<>G)_ + (*f*_CTT_ + *f*_GTT_ + *f*_CTC_ + *f*_GTC_…etc) × SFS_4F(C<>G)_) / (*f*_TTA_ + *f*_GTA_ + *f*_TTG_ + *f*_GTG_ + *f*_CTC_ + *f*_GTT_ + *f*_CTC_ + *f*_GTC_…etc).

Six parameters were estimated for each of the 75 non-synonymous site classes: the proportion of adaptive substitutions *α*, the rate of adaptive evolution relative to the mutation rate, *ω*_a_, the distribution of fitness effects for slightly deleterious mutations (DFE) modelled as a gamma distribution with the shape parameter *β* and the mean as the average selection strength against deleterious mutations *s*_d_, the average fitness effect of adaptive mutations, *s*_a_, as well as their proportion *λ*_a_. The demography parameters necessary as input into the DFE-alpha program were inferred from the synonymous SFSs, assuming a 3-epoch model, as implemented in DFE-alpha. The average segregating frequency of polymorphisms for each site class was calculated as *DAF* = (Σ_i_ iq_i_)/Σq_i_, where *q*_i_ represents the number of sites segregating at frequency *i* in the sample of sequences; 1 ≤ *i* ≤ 19, as we construct the SFSs from 20 sequences.

## Acknowledgements

JB was funded by the Austrian Science Fund (FWF, W1225-B20).

## Supplementary figures

**Figure S1.**
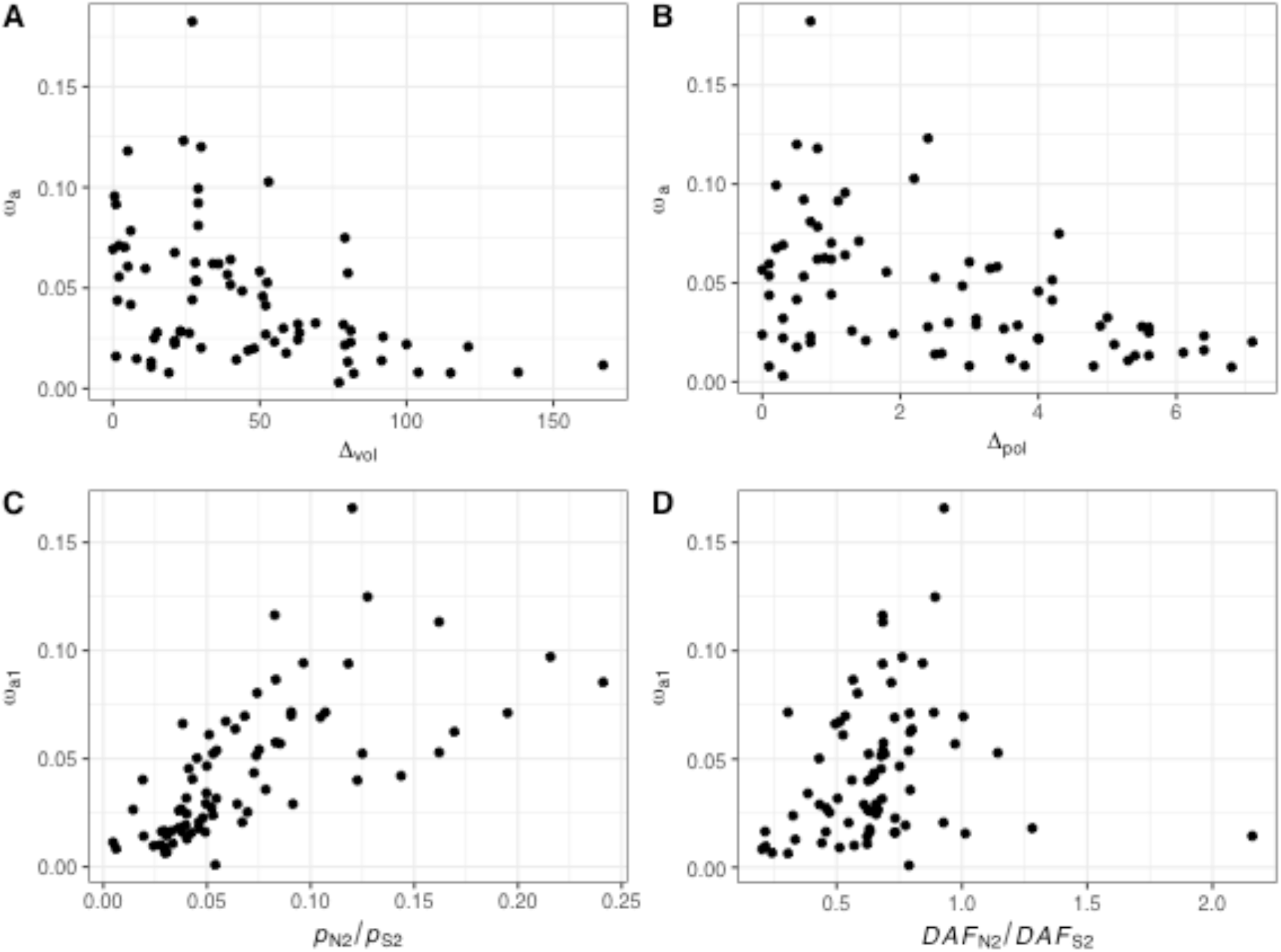
The X chromosome rate of adaptive evolution relative to the mutation rate (ω_a_) plotted against the difference in A) volume, B) polarity, C) *p*_N_/*p*_S_ and D) *DAF*_N_/*DAF*_S_.

**Figure S2.**
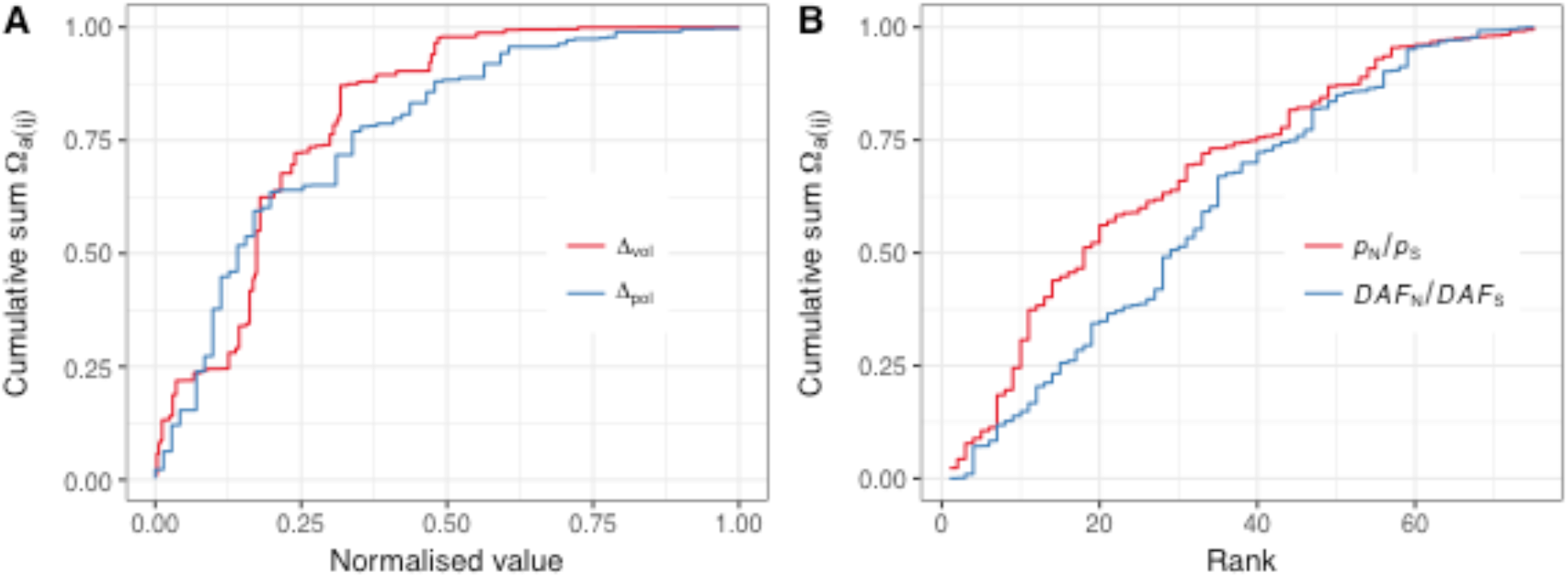
The cumulative number of adaptive substitutions on the X chromosome, Ω_a(ij)_, contributed by each pair of amino acids A) versus the normalised difference in volume and polarity, and B) versus the reverse rank of *p*_N_/*p*_S_ and *DAF*_N_/*DAF*_S_. The normalised difference in volume and polarity was calculated by subtracting the minimum difference, and then dividing by the maximum difference minus the minimum difference.

**Figure S3.**
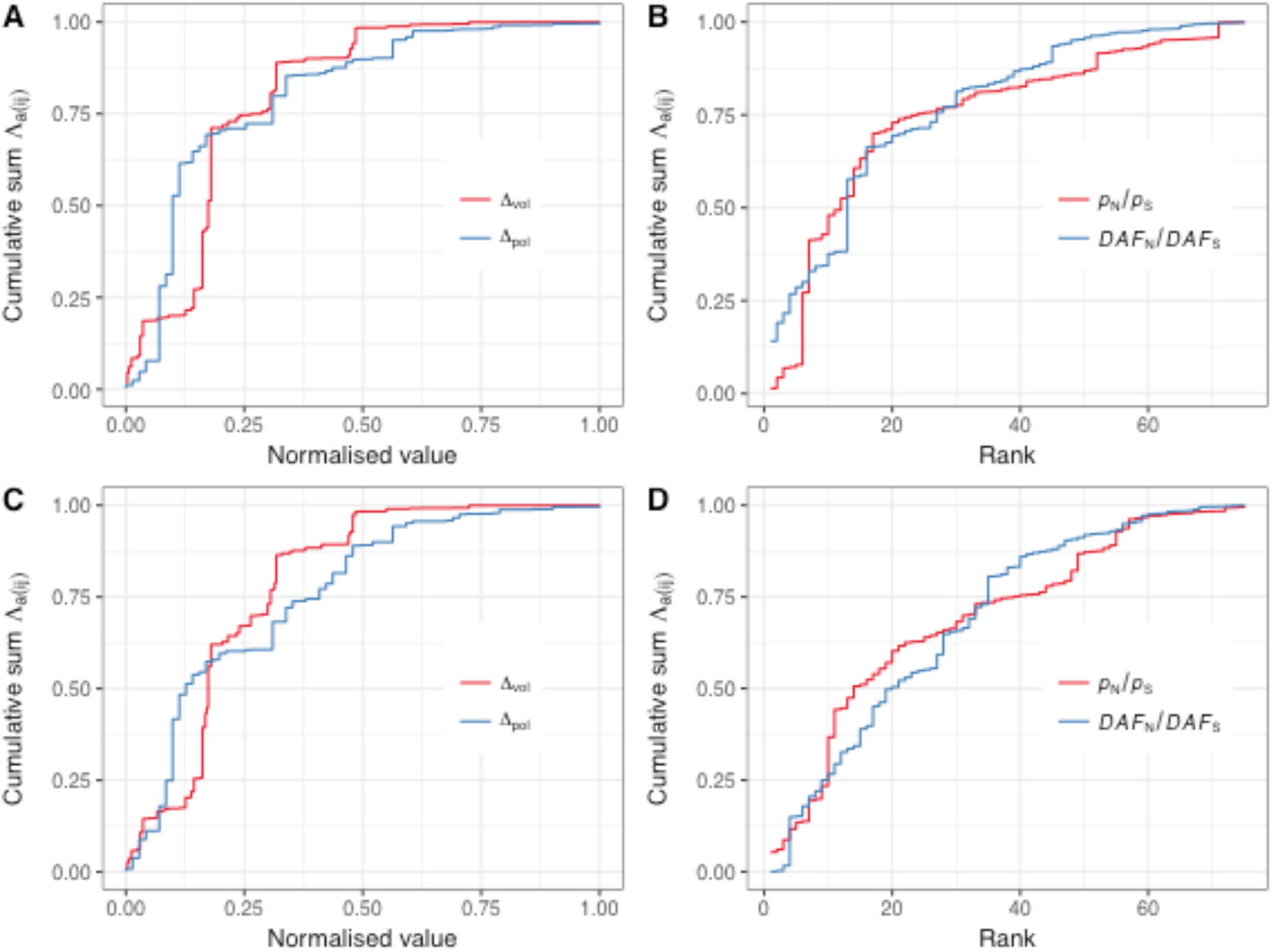
The cumulative number of adaptive mutations, Λ_a_, contributed by each pair of amino acids A) & C) versus the normalised difference in volume and polarity, and B) & D) versus the reverse rank of *p*_N_/*p*_S_ and *DAF_N_/DAF_S_*, for the autosomal data A) & B) and the X-chromosome C) & D). The normalised difference in volume and polarity was calculated by subtracting the minimum difference, and then dividing by the maximum difference minus the minimum difference.

**Figure S4.**
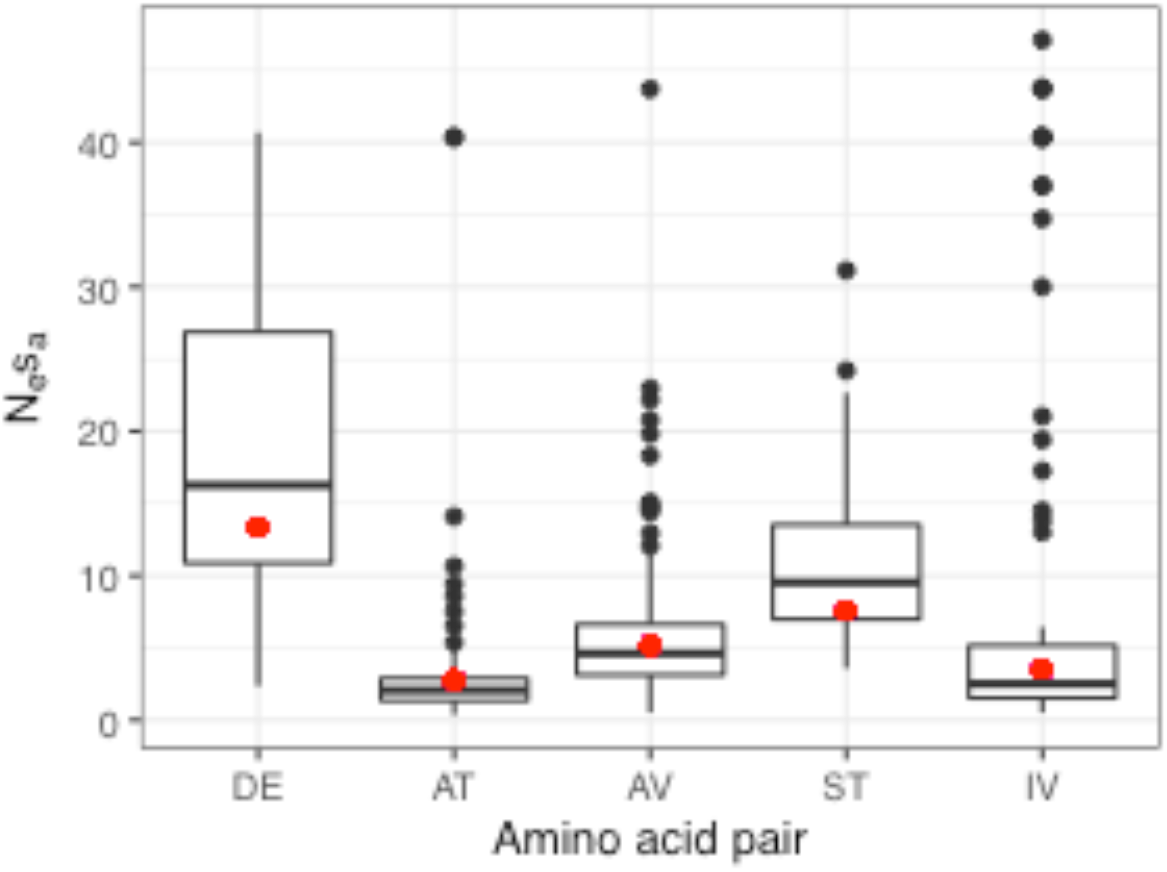
Distributions of the selection strength *N*_e_*s*_a_ for the five amino acid pairs with the most polymorphism data. Distributions were obtained by bootstrapping the SFSs of the amino acid pairs.

**Figure S5.**
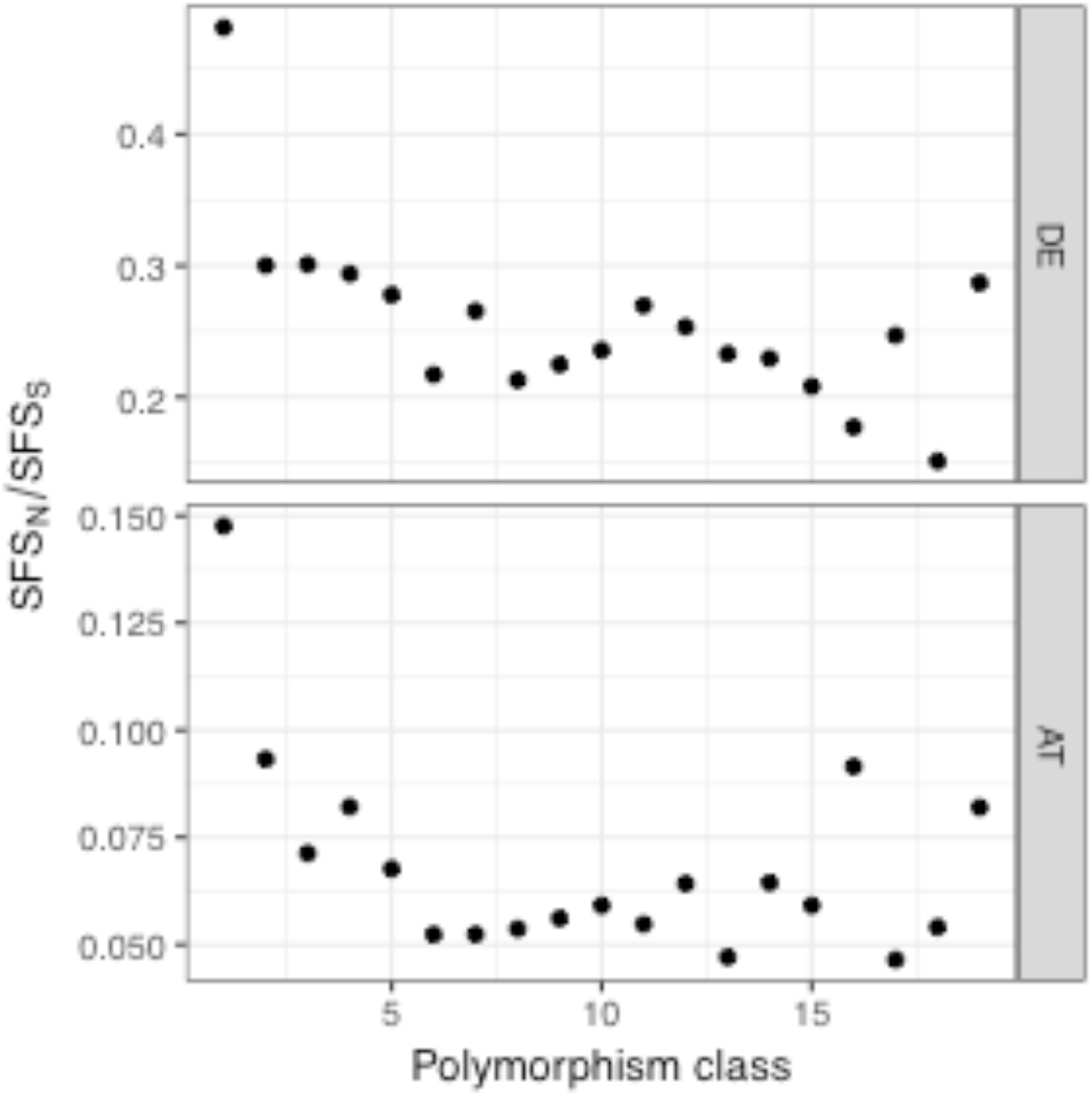
The ratios of the non-synonymous to the synonymous SFS (SFS_N_/SFS_S_) for the two amino acid pairs with the most polymorphism data.

**Figure S6.**
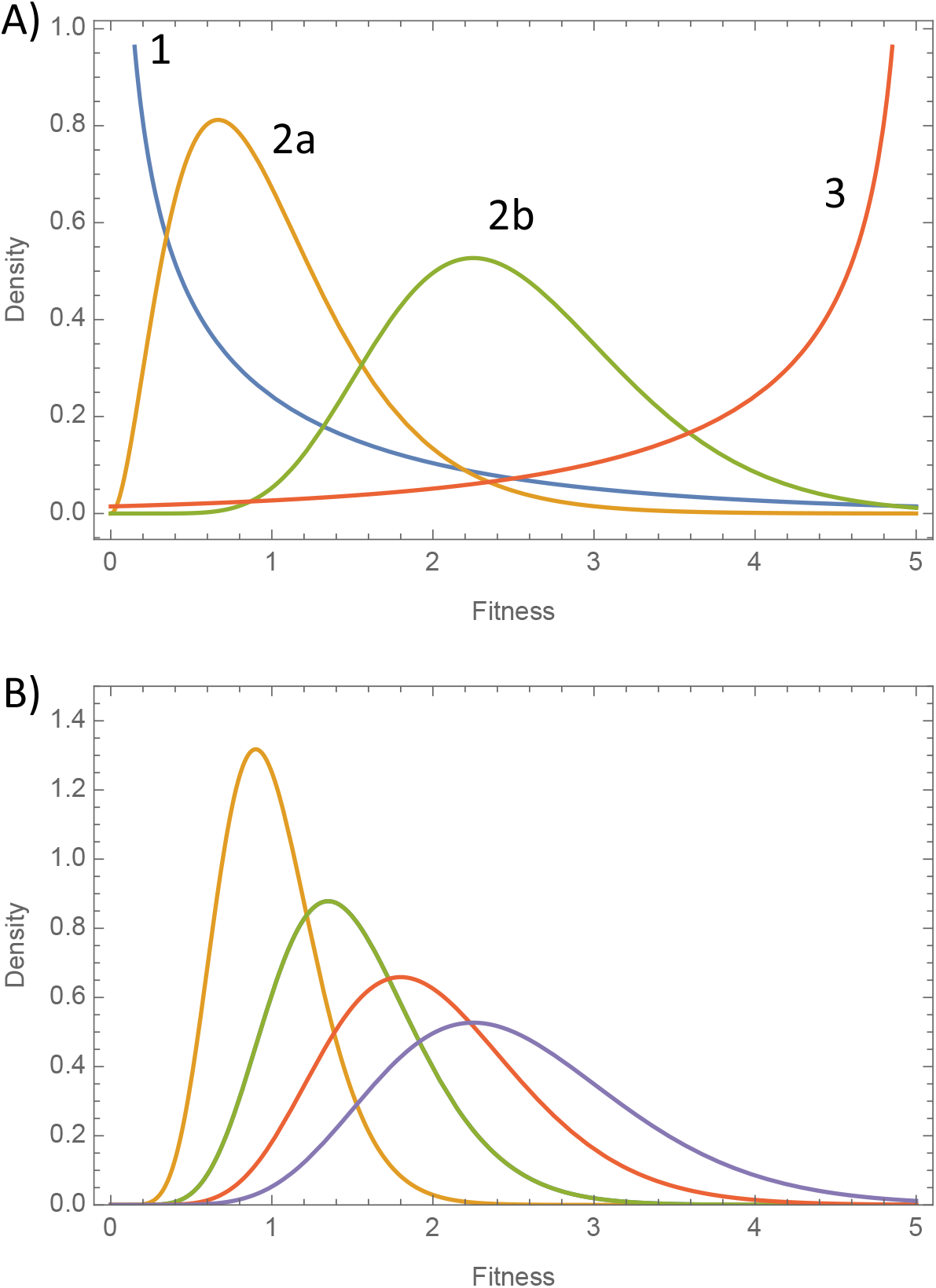
The underlying distribution of fitness effects amongst substitutions. Panel A shows the possible underlying distribution of effects amongst substitutions (or mutations) and panel B shows how distribution 2a in panel A could be composed of individual distributions each of which has a mean above the mode of the overall distribution. This would yield a negative relationship between the rate of adaptive substitution and a measure of fitness (e.g. the difference in polarity), even though the underlying distribution is such that mutations of intermediate effect size are the ones most commonly fixed.

